# PIAS1/PIAS4-Mediated SUMOylation of TDP-43 Is Induced by Oxidative Stress

**DOI:** 10.1101/2025.11.10.687649

**Authors:** Devon F. Pendlebury, Khloe Truong, Marie Heath, Jack C. Reidling, Partha Sarkar, Leslie M. Thompson

## Abstract

TAR DNA-binding protein 43 (TDP-43) is a conserved RNA and DNA binding protein that functions in transcriptional repression, pre-mRNA splicing and mRNA stabilization. Under pathological conditions found in multiple neurodegenerative diseases, TDP-43 shows aberrant mislocalization from the nucleus and cytoplasmic accumulation and aggregation. TDP-43 also appears to play a role in DNA damage repair, specifically in non-homologous end-joining (NHEJ), suggesting that nuclear depletion of TDP-43 may contribute towards the accumulation of DNA damage observed in TDP-43 proteinopathies. These TDP-43 pathological inclusions are decorated with post-translational modifications, most notably phosphorylation. SUMOylation (<u>S</u>mall <u>U</u>biquitin-like <u>Mo</u>difier) is a dynamic post-translational modification that regulates many protein properties and is implicated in neurodegenerative disease pathology. DNA damage repair proteins are commonly regulated through SUMOylation, and the SUMO E3 ligases PIAS1 and PIAS4 are required for efficient DNA repair of double-strand DNA breaks. Given these findings, we investigated TDP-43 SUMOylation and whether SUMO modification impacts TDP-43’s DNA damage repair function. We show that TDP-43 can be modified by SUMO1 and SUMO2/3 and confirm SUMOylation in response to oxidative stress. We also determine which regions of TDP-43 are SUMOylated and show that this modification is facilitated by the SUMO E3 ligases PIAS1 and PIAS4. Etoposide-induced DNA damage did not promote SUMOylation of TDP-43; studies are ongoing to determine the impact of TDP-43 SUMOylation on DNA repair.

## Introduction

TAR DNA-binding protein 43 (TDP-43) is a conserved RNA and DNA binding protein, involved in regulation of RNA metabolism including as a transcriptional repressor, in splicing regulation, and mRNA stabilization (*1-3*). TDP-43 normally localizes in the nucleus, but under pathological conditions is depleted from the nucleus and accumulates and aggregates in the cytoplasm. These aggregates are highly decorated with post-translational modifications, particularly phosphorylation and ubiquitination. TDP-43 pathological inclusions are found in numerous neurological diseases including amyotrophic lateral sclerosis (ALS), frontal temporal dementia (FTD), Alzheimer’s disease (AD), Huntington’s disease (HD), and limbic-predominant age-related encephalopathy (LATE) (*4-7*).

SUMOylation (<u>S</u>mall <u>U</u>biquitin-like <u>Mo</u>difier) is a dynamic post-translational modification which is a common property of proteins implicated in neurodegenerative disease pathology. SUMOylation can regulate many neuronal properties, including synaptic transmission, ion channel regulation, and neuronal development (*8*). Humans have five SUMO isoforms (*9-11*) with the stress-inducible forms, SUMO2 and 3, differing by only 3 amino acids and collectively referred to as SUMO2/3. SUMO is conjugated to lysine residues in target proteins through an enzymatic cascade similar to ubiquitination (*12*). SUMO proteins are initially translated as inactive precursors, activated with cleavage by the sentrin/SUMO-specific protease (SENP) family to expose a C – terminal di-glycine motif. Mature SUMO is activated by a single SUMO E1 activating enzyme, a heterodimer of SAE1 and 2, in an ATP dependent manner, then transferred to the sole E2 conjugating enzyme UBC9. Conjugation of SUMO to the target protein is facilitated by E3 SUMO ligases. Multiple SUMO E3 ligases have been identified, including the Protein Inhibitor of Activated STAT (PIAS) family and RANBP2 (*13, 14*). The E3 ligases also confer specificity to the target proteins (*12*). SUMO can then be removed from proteins through SENP activity. The balance of SUMO conjugation and deSUMOylation influences the properties of the target protein (*12*).

SUMOylation of TDP-43 has been previously reported, with studies suggesting TDP-43 is modified by SUMO2 primarily under stress induced by sodium arsenite (*15, 16*) and that TDP-43 SUMOylation is a) critical for regulating stress granule dynamics and b) may protect against aggregation. SUMO E3 ligases have been identified that can facilitate TDP-43 SUMOylation, although reports are not consistent across publications (*15-17*).

Multiple studies suggest that TDP-43 plays a role in DNA damage repair, including in non-homologous end-joining (NHEJ) mediated repair of double-strand breaks and the resolution of R-loops (*18-21*), suggesting that nuclear depletion of TDP-43 may contribute towards the accumulation of DNA damage seen in TDP-43 proteinopathies (*22-29*). DNA damage repair proteins are also commonly regulated through SUMOylation. For example, SUMOylation of replication protein A is essential for homologous recombination mediated repair (*30*), and SUMOylation of XRCC4 promotes its localization to the nucleus after DNA damage (*31*)(*19*). Additionally, the SUMO E3 ligases PIAS1 and PIAS4 are required for efficient DNA repair of double-strand DNA breaks (*32*). Given this data, we investigated whether TDP-43 is SUMOylated in response to DNA damage, which regions are SUMOylated and if TDP-43 SUMO modification modulates TDP-43’s DNA damage repair function. Here we show TDP-43 can be modified by SUMO1 and SUMO2/3 under basal conditions and in response to oxidative stress, but not in response to etoposide-induced DNA damage, which primarily created double-strand breaks. We identify sites that can be SUMO modified and that this modification is facilitated by the SUMO E3 ligases PIAS1 and PIAS4.

## Methods and Materials

### Immortalized cell culture

HeLa cells were cultured in Dulbecco’s Modified Eagle Medium (DMEM) supplemented with 10% fetal bovine serum (FBS) at 37°C, 5% CO_2_ and passaged at 95% confluency. SH-SY5Y cells were cultured in a 1:1 ration of minimum essential medium and Ham’s F12 Nutrient mix supplemented with 10% FBS. For experiments requiring transfection, HeLa were transfected with Lipofectamine 2000 (Invitrogen 11668027) and SH-SY5Y cells were transfected with Lipofectamine 3000 (Invitrogen L3000008).

### iPSC maintenance and neuronal differentiation

KOLF2.1 iPSCs with 19CAG repeats were differentiated and characterized as described by Smith-Geater *et. al*.(*33*).

### Plasmids

A pCMV-Myc-TDP-43 plasmid was a gift from Dr. Partha Sarkar, University of Texas Medical Branch (UTMB). TDP-43 domain cDNA constructs were purchased from GenScript and subcloned into pCDNA 3.1(+)N-Myc. His-SUMO1 and His-SUMO2 plasmids were a gift from Dr. Ron Hay, University of Dundee, UK. Human PIAS constructs were described as in (*34*).

### HeLa SUMOylation assay

HeLa cells were transfected with 1μg Myc-tagged proteins of interest and either: 1 μg vector, his-SUMO1 or his-SUMO2. Cells were harvested after 48 hours and lysed under denaturing condition (6M guanidine HCl, 100 mM NaH_2_PO_4_, and 10 mM Tris-HCl, pH 7.8). Lysates were sonicated 3x 10 seconds and 2.5% of lysate was set aside for input samples and purified via TCA precipitation. To isolate his-SUMO tagged protein, lysates were incubated with 20 μl magnetic cobalt bead slurry (Dynabeads His-Tag isolation and Pulldown, Invitrogen, Cat. No. 10104D) for 1 hour at room temperature. Beads were washed twice with wash buffer 1 (8M urea, 100 mM NaH_2_PO_4_, 10 mM Tris-HCl, pH 8), once with wash buffer 2 (8M urea, 100 mM NaH_2_PO_4_, 10 mM Tris-HCl, pH 6.3) and once with PBS. SUMO-conjugated proteins were eluted by resuspending beads in 1X LDS loading buffer, boiled for 10 min, and analyzed via sodium dodecyl sulfate polyacrylamide gel electrophoresis (SDS-PAGE) followed by western blotting.

### Analysis of endogenous SUMOylation

SH-SY5Y or iPSC derived striatal neurons were either untreated, treated with 250 µM sodium arsenite for 30 min, or 100 µM etoposide for 1 hour. Cell protein was then harvested and processed using Cytoskeleton’s Signal-Seeker SUMOylation 1 or 2/3 detection kits (BK165 and BK162). The kit protocol was followed using 500 µg protein.

### Western blotting

Protein samples were separated by SDS-PAGE using Invitrogen NuPAGE Bis-Tris Protein Gels, 4-12% with NuPAGE MOPS running buffer. Proteins were transferred onto a 0.45 µm nitrocellulose membrane (Bio-Rad #1620115). Near-infrared fluorescence of immunolabeled proteins was assessed using an Odyssey LICOR DLx. Protein quantification was calculated using Empiria Studio v2.0. Primary antibodies used were:

- Myc Tag Mouse clone 9E10 MilliporeSigma, 05419,1:1000,
- 6X-His Tag Polyclonal antibody, Inv
- itrogen, PA1983B, 1:1000
- TDP-43 Polyclonal antibody, Proteintech, 10782-2-AP, 1:1000
- Phospho-Histone H2A.X (Ser139) (20E3), Cell Signaling, 9718S, 1:1000
- Anti-SUMO-2/3 Antibody, clone 8A2, Millipore Sigma, MABS2039, 1:1000
- SUMO1 Antibody Mouse Monoclonal (Clone 5D8B16), Cytoskeleton, ASM01-S, 1:1000

### Immunofluorescence staining of SH-SY5Y cells

Cells were fixed with 4% paraformaldehyde for 10 min at room temperature, then washed three times with PBS. Cells were permeabilized with 0.1% Triton-X in PBS for 5 minutes and then blocked with 5% goat serum, 1% BSA, 0.1% Triton-X in PBS for 1 hour at room temperature. Coverslips were incubated in primary antibody diluted in blocking buffer overnight at 4°C (anti-myc-tag 9E10 1:1000 MilliporeSigma). After removing primary antibody, cells were washed three times with PBS and then incubated in secondary antibody diluted 1:1000 in blocking buffer for 1 hour at room temperature (Alexa Fluor 555). Cells were washed three times in PBS and then placed in PBS containing DAPI for ten minutes. Coverslips were mounted with Fluoromount-G and allowed to dry. 40x images were acquired on a Keyence BZ-X810 Widefield Microscope. 5 random images were taken per coverslip.

## Results

### TDP-43 can be SUMO modified by SUMO1 and SUMO2

TDP-43 is predicted to have multiple SUMO consensus sites (Fig 1A) (SUMOplot^TM^ Analysis Program and DeepSUMO(*35*) and has been shown previously to be SUMOylated (*15, 16, 36, 37*). To first confirm TDP-43 can be post-translationally modified by SUMO, we utilized a denaturing SUMOylation assay, using HeLa cells for their high levels of SUMOylation activity (*34, 38*). myc-TDP-43 and his-SUMO1 or 2 constructs were transfected into HeLa and after 48 hours his-SUMO-conjugated proteins were isolated using his-affinity beads under denaturing conditions to preserve only covalent SUMO modifications. An increase in TDP-43’s molecular weight is observed with both SUMO1 and SUMO2/3 his-affinity isolation (Fig1 B,C). Therefore, TDP-43 is subject to SUMOylation under basal conditions with addition of either exogenous SUMO1 or SUMO2/3.

**Fig 1.**
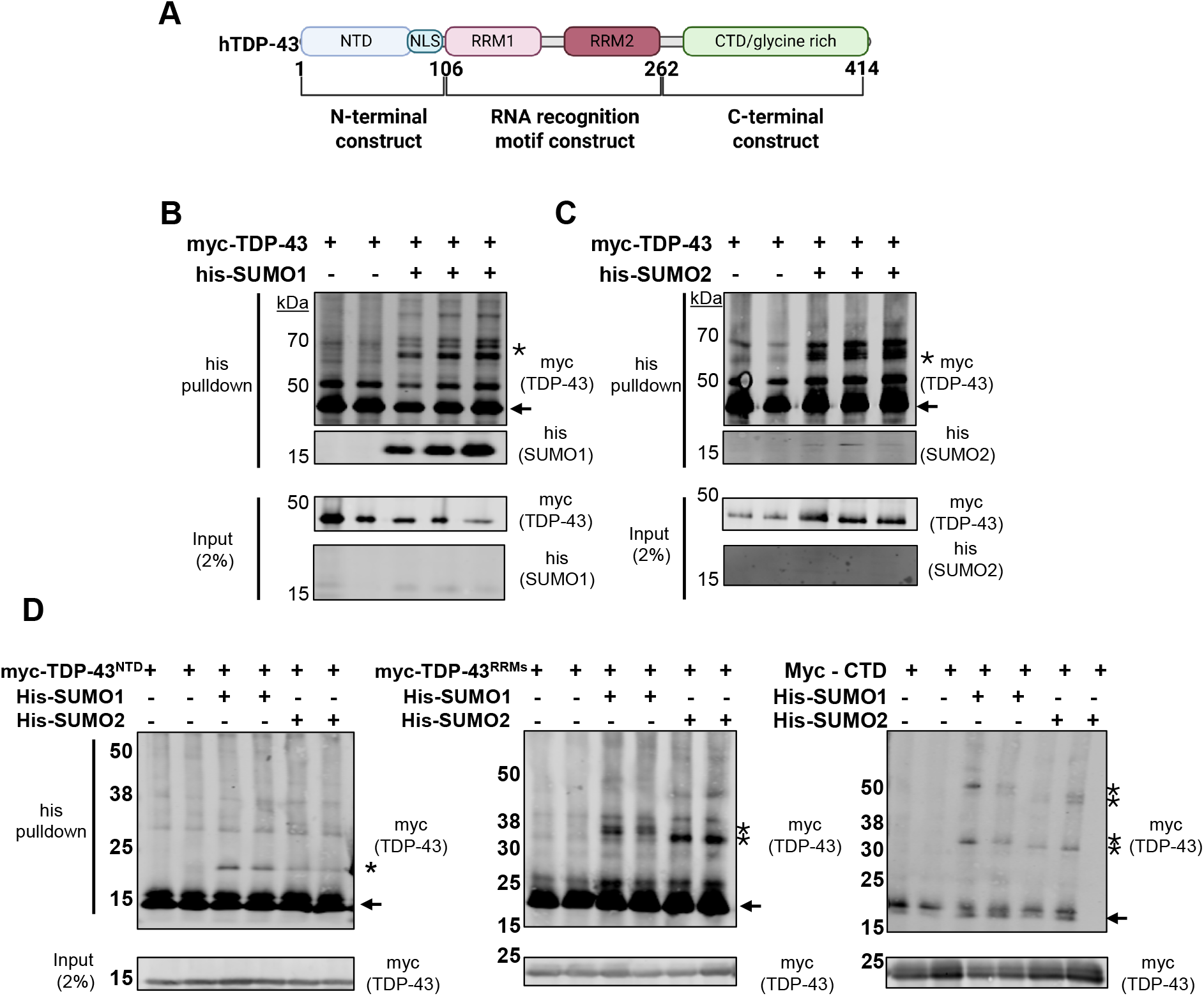
TDP-43 is modified by SUMO1 and SUMO2 across all domains. (A) Schematic representation of TDP-43 domains and the constructs used in SUMOylation assays. Created by Biorender https://BioRender.com/ay6vttk (B,C) Cobalt pulldowns under denaturing conditions from HeLa cells with his-SUMO1 (B) or his-SUMO2 (C) and myc-TDP-43. Immunoblots of input and pulldown samples are shown with arrows denoting the unmodified protein and asterisks denoting the SUMOylated protein throughout. (D,E,F) Cobalt pulldowns under denaturing conditions from HeLa cells of TDP-43^NTD^ (D), TDP-43^RRMs^ (E), and TDP-43^CTD^ with his-SUMO1 and his-SUMO2.

To better understand how SUMOylation might affect the properties and function of TDP-43 we investigated which specific regions of TDP-43 undergo SUMOylation. We repeated the denaturing SUMOylation assay with three myc-TDP-43 constructs (Fig 1A): TDP-43^NTD^ (aa 1-106) containing the N-terminal domain and nuclear localization signal (NLS), TDP-43^RRMs^ (aa 107-262) encompassing the RNA recognition motifs (RRMs), and TDP-43^CTD^ (aa 263-end) C-terminal domain (CTD) with the unstructured glycine-rich domain which drives aggregation in disease and harbors most disease-causing mutations (*39*). All three constructs showed shifts in molecular weight with SUMO1 and SUMO2 his-affinity isolation (Fig1 D,E,F), demonstrating SUMOylation occurs across all regions of the protein. These data suggest that SUMOylation may modulate multiple properties of TDP-43. Given the concentration of lysines in the NLS, SUMOylation in TDP-43^NTD^ could affect import into the nucleus. The RRM domains contain multiple lysines important for RNA binding and overall structure (*40*), and modifications in this domain could affect transcriptional and splicing regulation. Finally, TDP-43^CTD^ SUMOylation could potentially modulate TDP-43’s propensity to aggregate.

### PIAS1 and PIAS4 specifically facilitate the SUMOylation of TDP-43

We next sought to identify the SUMO E3 ligase responsible for TDP-43 SUMOylation, beginning with the PIAS family of E3 ligases given they are frequently responsible for the SUMOylation of proteins involved in DNA damage repair (*32, 41, 42*) and a global proteomics study showed that PIAS4 may SUMOylate TDP-43 (*17*). We carried out the same denaturing SUMOylation assay in HeLa cells, with limiting amounts of transfected his-SUMO2/3 to reduce baseline levels of TDP-43 SUMOylation, while also transfecting in either PIAS1, PIASxα, PIASxβ, PIAS3 or PIAS4. Both PIAS1 and PIAS4 overexpression resulted in increased TDP-43 SUMO2/3 modification (Fig 2), suggesting TDP-43 is a specific substrate of both, in agreement with recent studies (*15, 16*). None of the PIAS proteins facilitated modification of TDP-43 with SUMO1 (not shown). PIAS1 and PIAS4 are known to be key components of the DNA damage response (*32, 42*), play roles in both transcriptional regulation (*43, 44*), and stress response (*45, 46*), supporting a functional role in TDP-43 dynamics.

**Fig 2.**
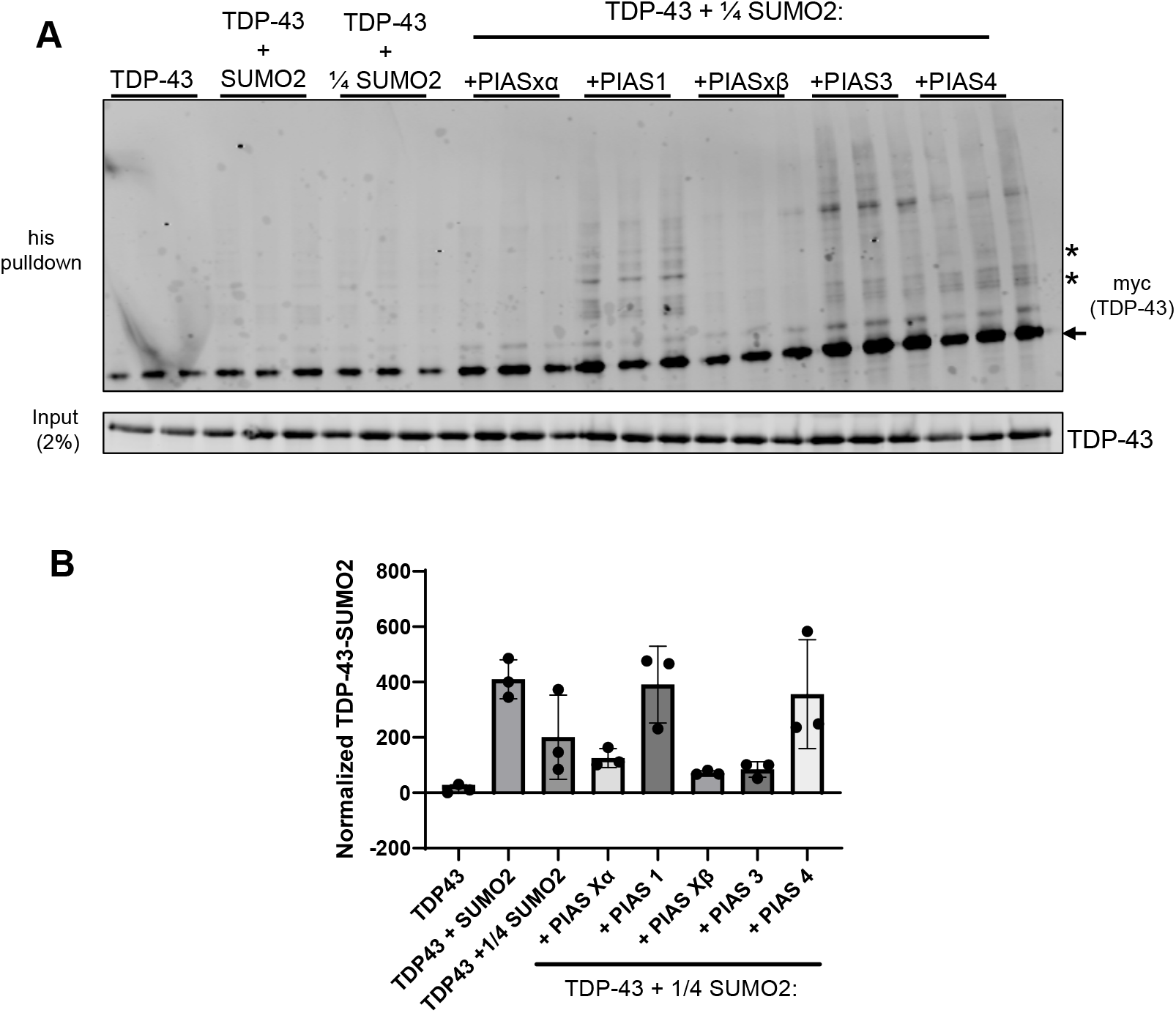
TDP-43 SUMO2-conjugation is facilitated by PIAS1 and PIAS4 E3 ligases. (A) Cobalt pulldowns under denaturing conditions of myc-TDP-43 with his-SUMO2 (1 µg plasmid) or limiting amounts of his-SUMO2 (0.25 µg plasmid). PIAS constructs were transfected to see if they could enhance SUMOylation when his-SUMO2 was limited. (B) The ratio of SUMO-modified to un-modified TDP-43 was calculated using Empiria Studio software by LICOR.

### TDP-43 is SUMOylated in response to oxidative stress but not DNA damage

To determine if TDP-43 is SUMOylated endogenously, without overexpression of SUMO or SUMO E3 ligases, we used Cytoskeleton’s Signal-Seeker^TM^ SUMOylation detection kits in SH-SY5Y neuroblastoma cells. No SUMOylation of TDP-43 with either SUMO1 or SUMO 2/3 was detected in either cell line under basal conditions (Fig 3A and B). Since SUMOylation with SUMO 2/3 often occurs to mitigate cellular stress (*47*), we treated both cell lines with sodium arsenite to induce oxidative stress (*48*). SUMOylation was seen in response to sodium arsenite stimulation (Fig 3A and B), in agreement with recent reports that TDP-43 SUMOylation is necessary for TDP-43 mobility in stress granules (*15, 16*). Given TDP-43’s role in DNA damage repair, we investigated whether TDP-43 was SUMOylated in response to DNA damage elicited by etoposide. Etoposide primarily induces double-strand DNA-breaks (*49*), which are the type of DNA damage TDP-43 has been reported to play a role in repair (*18-20*). Upon treatment with etoposide (*50*), indicated by expression of γH2AX (phosphorylated H2AX, a histone protein that acts as a sensitive marker for DNA double-strand breaks), neither cell type showed TDP-43 SUMOylation (Fig 3A and B), suggesting TDP-43 is not SUMOylated in response to this type of DNA damage. Further experiments will be needed to determine if other types of DNA damage induce SUMOylation of TDP-43, such as UV radiation single strand DNA damaging chemical reagents and reactive oxygen species.

**Fig 3.**
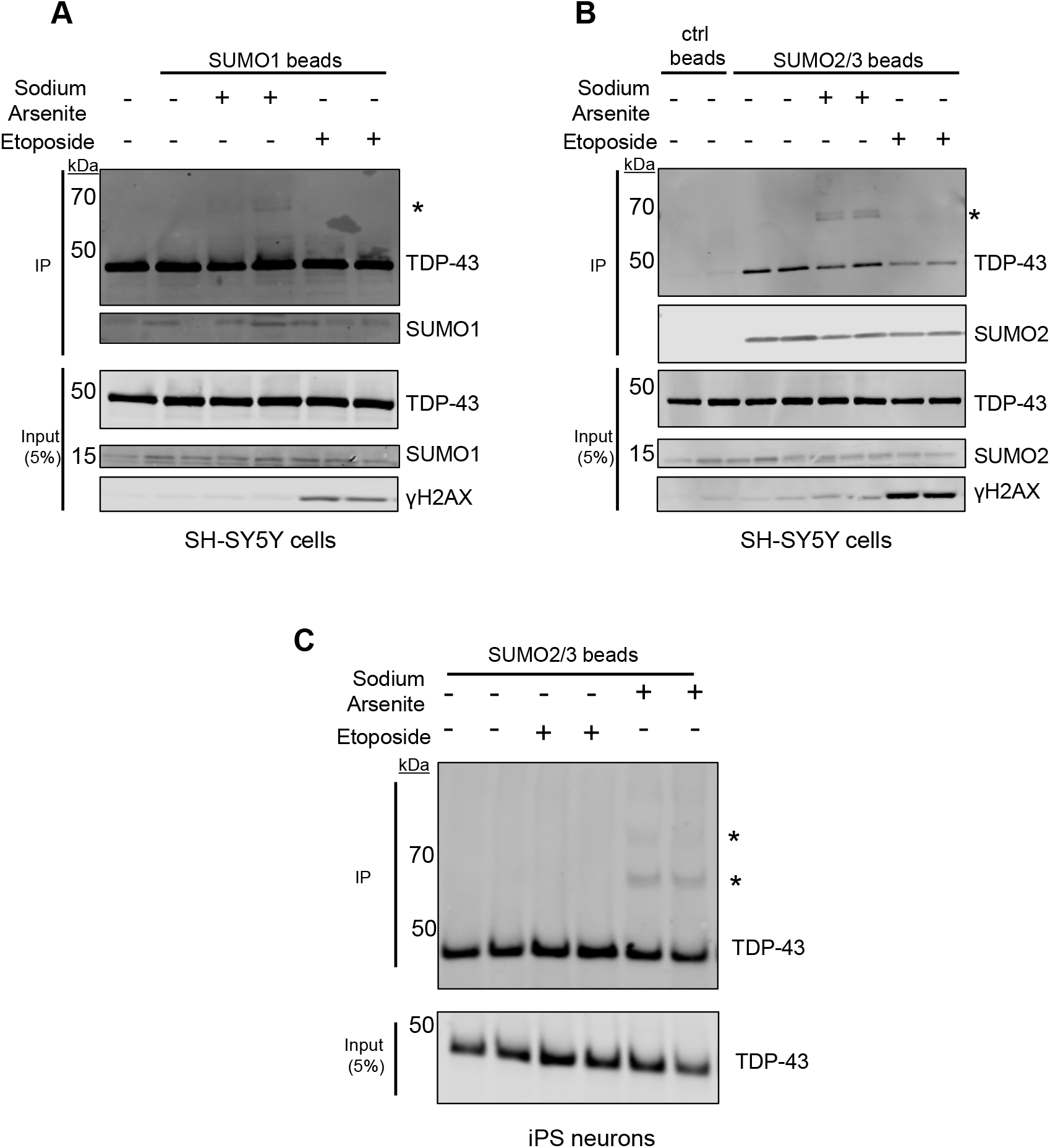
TDP-43 is SUMOylated in response to oxidative stress. (A, B) Immunoprecipitation of SUMO1(A) and SUMO2/3 (B) conjugated proteins from SH-SY5Y neuroblastoma cells using Signal-Seeker^TM^ SUMOylation Detection kits. Cells were either untreated, treated with 100 µM etoposide for 1 hour or 250 µM sodium arsenite for 30 min. (C) The same assay in (A,B) was carried out using induced pluripotent stem cell derived striatal neurons. Immunoblots from input (5% total protein) and pulldown samples are shown. Asterisks denote SUMOylated protein.

Since TDP-43 pathology is a common feature across numerous neurodegenerative diseases that affect neuronal subtypes, we confirmed that TDP-43 is also SUMOylated in neurons. We used induced pluripotent stem cell (iPSC) derived striatal neurons using the KOLF2.1J line, with the differentiation carried out as previously described (*33*). Results were consistent with the SH-SY5Y cells, with TDP-43 SUMOylation only detected after treatment with sodium arsenite (Fig 3C). Only SUMO2/3 modification was tested since TDP-43-SUMO2/3 conjugation was higher in SH-SY5Y cells.

### Mutation of TDP-43 SUMO sites reduces SUMOylation but does not affect subcellular localization

K136 is the highest confidence consensus SUMOylation site in TDP-43 and mutants have been previously evaluated (*36, 37*), although it was not directly tested whether a K136R mutation reduces SUMOylation. Additionally, K408 is the only lysine in the TDP-43^CTD^ construct to be within a predicted SUMOylation site. Therefore, we generated a TDP-43^K136/408R^ mutant (Fig 4A) and found that SUMOylation was reduced with SUMO2 (Fig 4B). To determine if TDP-43 localization changed in the SUMO-reduced construct, we transfected myc-tagged TDP-43^WT^, TDP-43^K136R^, TDP-43^K408R^, and TDP-43^K136/408R^ into HeLa cells given they are easily transfectable and used immunofluorescence to determine the localization of each construct. All constructs were primarily localized to the nucleus similar to wild-type TDP-43. The K136R mutation changed the staining from a diffuse nuclear signal to foci within the nucleus (Fig 4C). This is consistent with a previous study investigating the acetylation of K136 which found mutation of this residue formed nuclear foci/droplets (*51*). K136 interacts with nucleic acids (*40*) and mutation of this residue likely disrupts this binding, allowing unbound TDP-43 to aggregate or phase separate. This has also been observed in the presence of other TDP-43 mutations specifically designed to disrupt TDP-43 nucleic acid binding (*52-54*). These data suggest that SUMOylation at K136 may block binding to RNA and modulate TDP-43’s ability to regulate gene expression and splicing.

**Fig 4.**
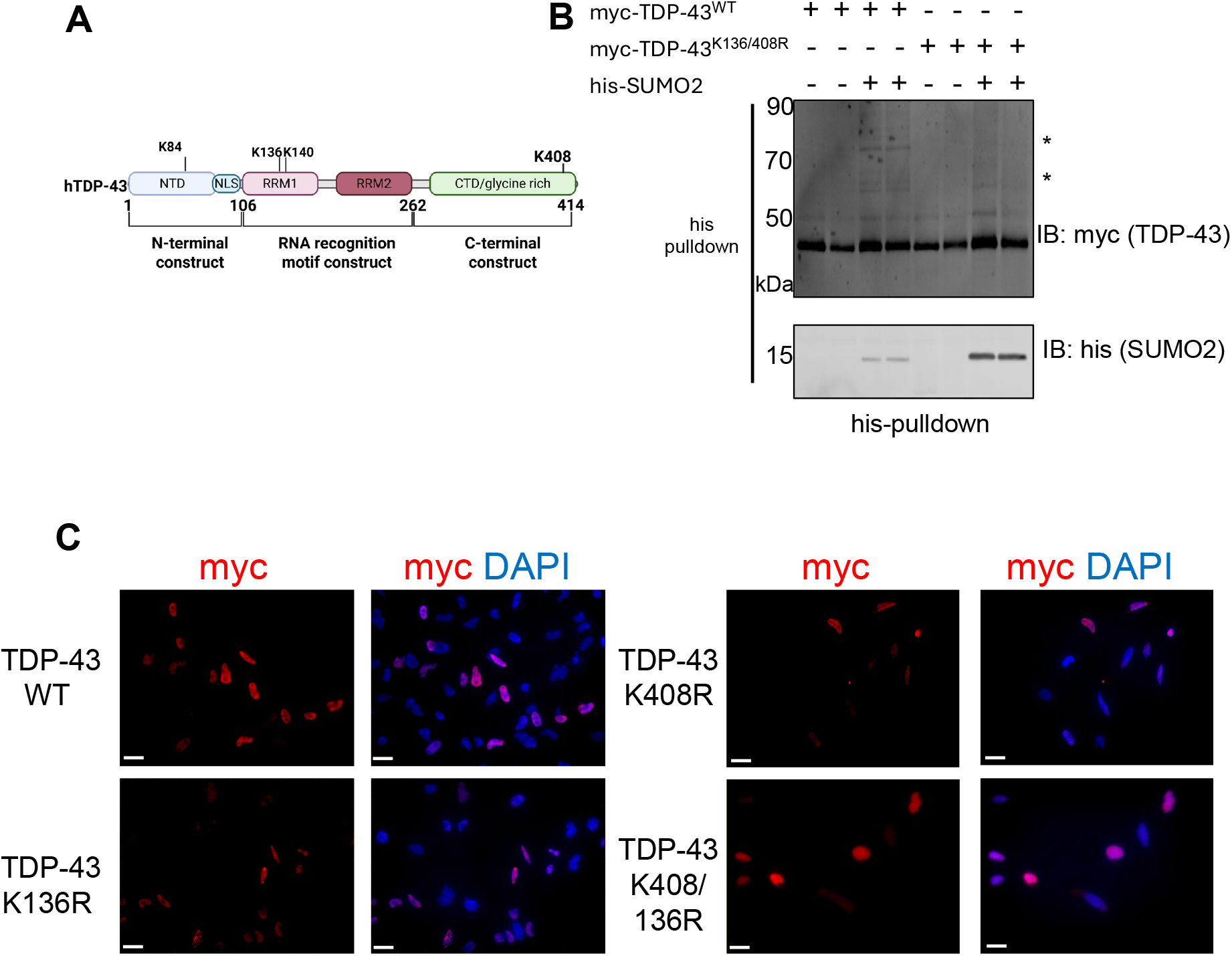
Mutation of K136 and K408 reduces TDP-43 SUMOylation. (A) Schematic of TDP-43 with predicted SUMOylation sites marked, as determined using SUMOplot^TM^ Analysis Program and DeepSUMO(*27*). Created by BioRender Created in BioRender https://BioRender.com/szete9f (B) Cobalt pulldown under denaturing conditions with my-tagged TDP-43^WT^ or TDP-43^K136/408R^ with his-SUMO2. Immunoblots from input (5% total protein) and pulldown samples are shown. Asterisks denote SUMOylated protein. (C) Representative fluorescent microscopy images of myc-tagged TDP-43 constructs in SH-SY5Y cells. Scale bar 5µm.

To determine if SUMOylation alters the subcellular localization of TDP-43, we overexpressed PIAS1 or PIAS4 in HeLa cells and monitored TDP-43 localization by immunofluorescence. TDP-43 remained nuclear under all conditions, suggesting excess SUMOylation does not drive TDP-43 from the nucleus to the cytoplasm (data not shown).

## Discussion

TDP-43 is depleted from the nucleus and accumulates in cytoplasmic inclusions in multiple neurodegenerative diseases including ALS, FTD, HD and AD (*4-7*). These inclusions are post-translationally modified, particularly by phosphorylation and ubiquitination, but other less studied modifications of TDP-43, including SUMOylation, acetylation, and citrullination, also occur (*15, 16, 36, 37, 51, 55-57*). Here we show that TDP-43 can be SUMO-modified by exogenous SUMO1 and SUMO2/3. We also report that TDP-43 SUMOylation is enhanced by SUMO E3 ligases PIAS1 and PIAS4 in response to oxidative stress but not by etoposide-induced double-strand DNA damage, in both immortalized cancer cell lines and iPSC-derived neurons. Given the increase in cellular stress in neurodegenerative disorders, as well as in aging neurons, it is likely that there is an increase in TDP-43 SUMOylation in these populations. TDP-43 SUMOylation could be a response to prevent TDP-43 aggregation in the cytoplasm. Future studies examining if TDP-43 aggregates are SUMOylated in patient tissues are needed, as well as more in-depth research into whether SUMOylation of TDP-43 in disease is protective or pathogenic and the TDP-43 functions that may be modulated by this PTM. Recent results show that there is a correlation between motor neurons with decreased cytoplasmic PIAS4 levels and TDP43 inclusions (*15*) suggesting SUMOylation is beneficial in preventing aggregation.

SUMOylation of pathogenic proteins is a common feature of neurological diseases (*58*). In HD, both WT and mutant Huntingtin protein (HTT) can be SUMOylated, which is enhanced by PIAS1, resulting in the accumulation of insoluble mutant HTT in cell culture (*34*). In human HD postmortem striata, an increase in insoluble SUMO-conjugated proteins in addition to HTT is also observed (*34, 59, 60*). Knockdown of PIAS1 improves numerous HD phenotypes including behavioral and motor phenotypes in HD mice, reduction of HTT inclusions, increased genomic integrity, correction of transcription dysregulation and rescue of neuronal activity (*42, 59, 60*). Additionally, α-synuclein, which aggregates in Parkinson’s disease, can be SUMO modified with mixed reports on whether this drives or inhibits aggregate formation (*61-63*). Tau, which accumulates in AD, can also be SUMOylated which increases its aggregation (*64, 65*). Other proteins which are involved in ALS pathogenesis including superoxide dismutase 1 (SOD1) and fused in sarcoma/translocated in liposarcoma (FUS/TLS) have also been reported to be SUMOylated (*66-69*). The role of SUMOylation on FUS pathology has not been explored but SUMOylation of SOD1 promotes its aggregation. Together these data support a key role for SUMOylation in neurodegenerative diseases. Given the number of TDP-43 proteinopathies, further studies on TDP-43 SUMOylation are critical. Cumulatively, these data emphasize that manipulation of the SUMO pathway is a promising therapeutic avenue for numerous neurodegenerative diseases.

Our results are in agreement with a previous study which reported that PIAS4 mediates TDP-43 SUMOylation in response to oxidative stress, and that this property is important for TDP-43 mobility and solubility in stress granules (*15*). Similarly, another report suggests mutation of K408 impairs cellular stress in neurons (*16*), further highlighting the importance of TDP-43 SUMOylation in oxidative stress responses and the formation of stress granules.

We did not observe any change in TDP-43’s SUMOylation status based on induction of DNA damage with etoposide and monitored with γH2AX, suggesting that SUMOylation of TDP-43 is not induced by DNA damage, although it may still play a role in DNA repair. Perhaps given its nuclear localization, modifications are not needed to recruit TDP-43 to sites of DNA damage, or possibly it is recruited by interacting with other SUMOylated DNA repair proteins through its predicted sumo-interaction motifs (DeepSUMO(*35*)). It will be of interest to determine whether cytosolic mislocalization alters the response to SUMO modification in influencing DNA damage responses. It is also possible that TDP-43 may not play a role in etoposide induced DNA damage repair, and other methods of DNA damage induction need to be tested. Additionally, even if TDP-43 is not directly SUMOylated in response to DNA damage in our assays, it may just be that only a small subset of TDP-43 in the cell is involved in DNA damage repair, while the rest of the population remains involved in RNA regulation. As DNA damage repair complexes often interact via interactions between SUMO and SUMO-interaction motifs, it would be interesting to see if these mutants affect the interactions of TDP-43 with other DNA repair proteins.

Our data shows that modulation of SUMOylation does not alter TDP-43’s subcellular localization in immortalized cells, either by mutants which reduce SUMOylation or overexpressing PIAS1 and PIAS4 to promote SUMOylation. Further studies are needed to determine if SUMOylation affects the aggregation and activity of TDP-43 when it is already in the cytoplasm, such as in ALS, and whether there are differential effects when evaluated in neurons. Although our TDP-43^K136/408R^ construct showed reduced SUMOylation, care must be taken when interpreting these results, as lysines can frequently modified by multiple post-translational modifications. K136 in TDP-43 can also be acetylated (*51*) and ubiquitin can often compete with SUMO for the same residues. Our K136R mutant showed less disperse nuclear localization and formed nuclear foci, consistent with previous publications studying this residue (*51*) and other mutations which disrupt nucleic acid binding (*52-54*). Our experiments depended on the overexpression of proteins using plasmids, which may not represent endogenous phenotypes.

In summary, we show that TDP-43 SUMOylation is enhanced by the E3 ligases PIAS1 and PIAS4 in response to oxidative stress, as previously shown, but not in response to DNA damage. Mutation of K136/408R reduced SUMOylation but did not have an effect on TDP-43 subcellular localization. Overexpression of E3 ligases also did not alter its subcellular localization, however we do not yet know how SUMOylation affects cytoplasmic TDP-43 localization. Further studies are needed to determine TDP-43 SUMOylation is altered in TDP-43 proteinopathies *in vivo* and if its modulation is therapeutically beneficial.

## Acknowledgements

We would like to thank the Thompson lab for critical feedback. This work was supported by the following National Institutes of Health (NIH) grants: R35NS116872 (L.M.T.) and F32NS136263 (D.F.P.) and Huntington’s Disease Foundation fellowship (D.F.P.).

